# A single QTL with large effect is associated with female functional virginity in an asexual parasitoid wasp

**DOI:** 10.1101/2020.11.05.370544

**Authors:** Wen-Juan Ma, Bart A. Pannebakker, Xuan Li, Elzemiek Geuverink, Seyed Yahya Anvar, Paris Veltsos, Tanja Schwander, Louis van de Zande, Leo W. Beukeboom

## Abstract

During the transition from sexual to asexual reproduction, a suite of reproduction-related sexual traits become superfluous, and may be selected against if costly. Female functional virginity refers to asexual females resisting to mate or not fertilizing eggs after mating. These traits appear to be among the first that evolve during the gradual transition from sexual to asexual reproduction. The genetic basis of female functional virginity remains elusive. Previously, we reported that female functional virginity segregates as a single recessive locus in the asexual parasitoid wasp *Asobara japonica*. Here, we investigate the genetic basis of this trait by quantitative trait loci (QTL) mapping and candidate gene analyses. Consistent with the segregation of phenotypes, a single QTL of large effect was found spanning over 4.23 Mb and comprising at least 131 protein-coding genes, of which 15 featured sex-biased expression in the related sexual *Asobara tabida*. We speculate that two of these 15 genes may be of particular interest: *CD151 antigen* and *nuclear pore complex protein Nup50*. Overall, our results are consistent with a single gene or a cluster of linked genes underlying rapid evolution of female functional virginity in the transition to asexuality. Once a mutation for rejection to mate has swept through a population, the region comprising the gene(s) does not get smaller due to lack of recombination in asexuals.

## Introduction

A suite of reproduction-related sexual traits become superfluous during the transition from sexual to asexual reproduction (e.g. Jeong & Stouthamer 2005; Pannebakker *et al*. 2005; Kraaijeveld & Vavre 2009; Russell & Stouthamer 2011; Carson *et al*. 2013; Schwander et al. 2013; Ma *et al*. 2014a; Kraaijeveld *et al*. 2016; Parker *et al*. 2019). Asexual females do not require mating with males for offspring production, and produce only daughters. As a consequence, costly sexual traits are expected to be selected against in asexual lineages (Ma, Vavre, & Beukeboom, 2014; van der Kooi & Schwander, 2014). Consistent with this expectation, there is good evidence for rapid reduction of sexual traits in asexual females, especially of traits linked to mate attraction and copulation, which are believed to be particularly costly in many species (Schwander et al., 2013; van der Kooi & Schwander, 2014).

Traits reducing mate attraction or copulation probability are of particular interest in hymenopteran insects, where they have been referred to as “female functional virginity” (Jeong & Stouthamer, 2005; Russell & Stouthamer, 2011; Stouthamer, Russell, Vavre, & Nunney, 2010). In hymenopterans, virgin sexual females produce only sons because sex determination relies on haplo-diploidy (males develop from unfertilized, haploid eggs, females from diploid eggs) (Whiting, 1933). As a consequence, female functional virginity can be favored during the spread of facultative asexuality in a sexual population, because it favors the production of males in female-biased populations (Huigens & Stouthamer, 2003).

Reduced sexual traits associated with female functional virginity have been reported in at least five different asexual hymenopteran species (Pijls et al. 1996; Jeong & Stouthamer, 2005; Pannebakker *et al*., 2005; Russell & Stouthamer, 2011; Stouthamer, Russell, Vavre, & Nunney, 2010; Ma *et al*., 2014a). The genetic and molecular basis underlying female functional virginity remains elusive, which is in part due to the inability to perform crosses in asexual organisms. However, crosses are possible in some hymenopteran asexual species, because asexuality is caused by infection with maternally transmitted endosymbionts, such as *Wolbachia*. When asexual females are cured from their endosymbionts via antibiotic treatment, they produce males and these males can be mated to females from related sexual strains (Pannebakker *et al*., 2005; Jeong & Stouthamer, 2005; Russell & Stouthamer, 2011; Ma *et al*., 2014a). The current evidence of the genetic mechanisms resulting in female functional virginity in asexual hymenopterans points to mutations in few genes with relatively strong phenotypic effects (reviewed in van der Kooi & Schwander, 2014), although these results may also be caused by the Beavis effect, which postulates an overestimation of QTL effect sizes when sample size is small (Beavis, 1994).

*Asobara japonica* is a parasitic wasp with both sexual and all-female asexual populations that are geographically separated. Asexual populations are distributed across mainland Japan, whereas sexual populations occur on southern islands (Murata *et al*., 2009). *A. japonica*, like all hymenopterans, has haplodiploid reproduction (Werren, 1997; Werren *et al*., 2008; Mateo Leach *et al*., 2009; Giorgini *et al*., 2010; Kageyama *et al*., 2012; Ma *et al*., 2014). Asexuality in *A. japonica* is induced by the bacterial endosymbiont *Wolbachia*, and unfertilized haploid eggs laid by endosymbiont-infected females typically undergo diploidization and develop as females (Kremer *et al*., 2009; Reumer *et al*., 2012). Removing the infection by antibiotic treatment results in the production of haploid males from unfertilized eggs. Asexual females also occasionally produce haploid males under natural conditions (Heath *et al*., 1999; Reumer *et al*., 2012). Although these males usually do not have any mating opportunities in nature, in the laboratory they can be mated with females of sexual strains.

A previous study in *A. japonica* revealed that asexual females are less attractive than sexual females, and completely refuse to mate (Ma *et al*., 2014). Female functional virginity in asexual *A. japonica* is thus likely governed by genes mediating female mating propensity. Introgression of alleles from an asexual strain into a sexual one for four consecutive generations revealed that a single recessive locus with major effects caused introgressed females to produce only haploid sons, suggesting a relatively simple genetic architecture for female functional virginity (Ma *et al*. 2014a).

In the current study we identified the genomic regions and possible candidate genes associated with female functional virginity in *A. japonica*. We genotyped the sex-asex introgressed females that were previously analyzed for resistance to mating and produce only haploid sons (Ma *et al*. 2014a). We constructed a genetic map, a genome assembly of *A. japonica* from PacBio long reads sequencing, and subsequently performed a quantitative trait loci (QTL) analysis. In addition, Illumina short-reads of sexual and asexual strains of *A. japonica*, and a transcriptome dataset of the sexual species *Asobara tabida*, were used to infer candidate genes.

## Methods

### Wasp strains and culturing

A sexual and a *Wolbachia*-infected, asexual strain of *Asobara japonica* were used. The sexual strain originated from the Amami-Oshima island (AO) and the asexual strain from Kagoshima (KG), both in Japan, and have been cultured in the laboratory since 2009. These two strains are closely related (Murata *et al*., 2009; Reumer *et al*., 2012), which minimizes the probability of genetic incompatibility in crosses. *A. japonica* was cultured on second-instar *Drosophila melanogaster* larvae as hosts at 25°C, with a 16L: 8D light-dark cycle and 60% relative humidity (for details see Ma et al. 2013, 2014).

Sons from asexual females (so called “asexual males”) were obtained by antibiotic treatment of *Wolbachia*-infected females (see below) or directly collected from mass cultures in which males occur at low frequency (i.e. ~0.7%, N=6000 individuals). As spontaneously occurring males are sometimes diploid (Ma *et al*., 2015), flow cytometry was used to ensure that all males used in experiments were haploid (for details see Ma *et al*., 2013).

Antibiotic treatment (to obtain “asexual” males) was performed by providing *Wolbachia*-infected females with *Drosophila* hosts cultured on food containing10 mg of rifampicin for 1g yeast powder (see Ma *et al*. 2014a). Rifampicin reduces the *Wolbachia* titer, but has little impact on development in *Asobara* (Dedeine et al., 2001). Complete removal of *Wolbachia* was confirmed by the production of only male offspring by the treated females.

### Introgression cross design

To investigate the genetic basis of female functional virginity, measured as females producing haploid sons only, asexual KG males were mated to virgin sexual AO females in the first generation, and then to introgressed F1 females for three consecutive generations (details in Ma *et al*. 2014a, Figure 1b). Briefly, virgin females were collected by individually isolating wasp pupae in plastic vials (diameter 2.4 cm, height 7.5 cm) containing a layer of agar to control humidity (Ma *et al*., 2013). These virgin females were individually paired with a male from the asexual strain for 24 hours, and subsequently offered approximately 100 second-instar *D. melanogaster* larvae for oviposition for 36 hours. The resulting wasp pupae were isolated from parasitized hosts to prevent mating upon emergence. Females emerging from these pupae (the F1 generation) were collected and backcrossed to males of the asexual KG strain, by individual pairing, to produce the offspring of the next generation (Ma *et al*. 2013, 2014).

**FIG. 1.**
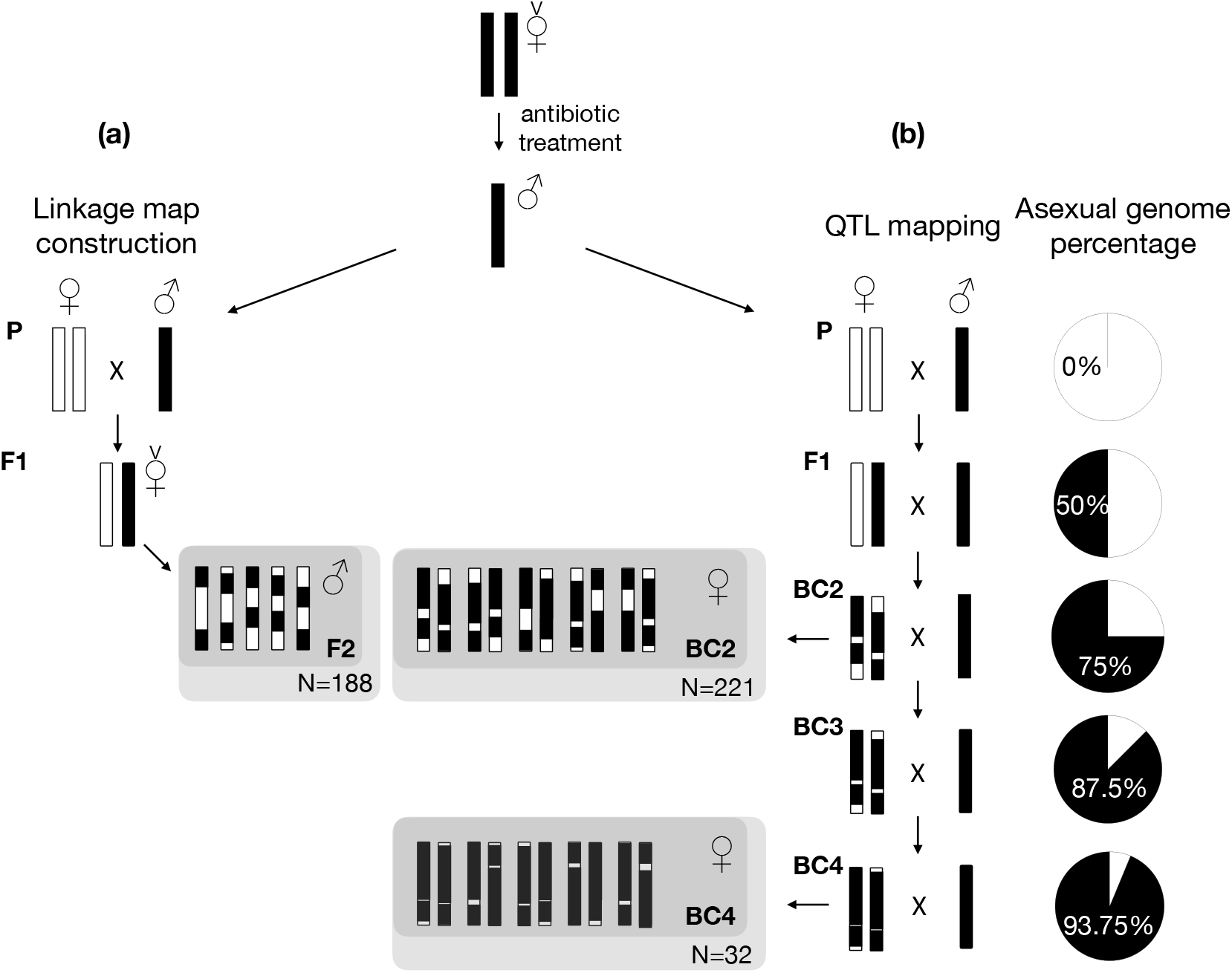
Cross schemes used to generate F2 recombinant males for linkage map construction (a), and four generations of backcrosses for QTL mapping of the female functional virginity trait (b). The AO sexual female (white) is crossed to the KG asexual male (black) to generate F1 females that were set up as virgins to produce recombinant F2 males. The F2 males (a) in grey shade were used for genotyping and linkage map construction, and females from backcross generation 2 (BC2) and 4 (BC4) (b) in grey shade were used for determining female mating success and QTL mapping. Black and white mosaic bars denote relative component of sexual and asexual genomes in consecutive generations. The pie chart shows the percentage of asexual genome component (in black) in diploid females in consecutive introgressed generations (modified from Ma *et al*. 2014a).

Under this cross design, the proportion of asexual alleles in females of each generation is expected to increase from 50% in F1, to 75% in BC2, 87.5% in BC3 and a 93.8% in BC4 (Ma *et al*. 2014a; Figure 1b). For each backcross, the emerging wasps were anaesthetized with CO_2_, counted and sexed (Ma *et al*. 2013, 2014). For each backcross generation, a subset of introgressed females (N=49-225 depending on the generation) was sampled from the introgression experiment to determine their offspring sex ratio, by pairing a single female with a sexual male for 24 hr and allowing 36 hr for oviposition. Each female was scored for production of exclusively male or male and female offspring. The thus obtained binomial trait was used as the phenotype for QTL mapping.

### Genome-wide SNP markers development

To enable genetic analysis, SNP markers were generated from Illumina short reads of 30-pooled individuals of both the sexual AO and asexual KG strains (BioProject: PRJNA661661, accession number: SRR12618696-SRR12618699). Genomic DNA was extracted with a modified high-salt protocol (adjusted from Aljanabi, 1997, see details in the next section). DNA libraries were prepared at the University Medical Center Groningen Sequencing Facility and sequenced on an Illumina HiSeq2000. SNP markers were developed from the Illumina short reads using the *de novo* genome-wide DIAL (De novo Identification of ALleles) computational pipeline, which identifies SNPs between two closely related genomes without needing a reference genome (Ratan *et al*., 2010). To prevent long CPU times, only a subsample of the reads (1,172.5Mb or 3.9 - 4.2x coverage of the estimated wasp genome of 282-304 Mb, see K-mer analysis below) were analyzed. The DIAL pipeline first removes duplicate reads and builds a short consensus sequence (ranging from 95-300 bp), which it then aligns to reads from previous runs to form clusters. Clusters with few nucleotide differences are selected and putative SNPs are detected and filtered based on the read quality at the variant position (minimum Phred-quality score used was 20).

The DIAL pipeline initially generated a database of 21,976 SNPs with their flanking region sequences and information about coverage depth and phred-quality scores for each of the two strains. The SNP coverage ratio R is defined as the DIAL-generated coverage depth (d_SNP) of the SNP nucleotide, divided by the average coverage depth (d_reads) of the selected reads (against the estimated genome size): R=d_SNP/d_reads. Following author recommendations (Ratan *et al*., 2010), only markers with R > 1.5 were selected for further verification (n=10,001). Next, minimum coverage 3 and minimum base pair quality score 30 were used for further filtering which resulted in 951 remaining SNPs. Lastly, mitochondrial, bacterial and human homologous fragments were removed from the candidate SNP cluster list based on the BLAST results from NCBI sequence database, yielding 420 usable SNPs of which 130 were randomly selected for linkage map construction.

A subset of these 130 SNPs (57) was tested for amplification in both strains with primer pairs designed with Perlprimer (version 1.1.21, Marshall, 2004, Table S1) and Sanger sequenced (see below). This yielded 47 confirmed SNPs, to which another 66 randomly selected SNPs from the filtered SNP pool were added, for a total of 113 SNPs to be used for in the linkage map and QTL analyses.

The Illumina short reads (KG and AO strains) were also used for estimating the genome size for each strain of *A. japonica* (KG: 281~304 Mb; AO: 280~284 Mb). The genome size was estimated by fitting the k-mer distribution of the Illumina short reads, obtained from Jellyfish v2.1.0 (Marçais & Kingsford, 2011), using GenomeScope v2 (http://qb.cshl.edu/genomescope/genomescope2.0/) (Vurture *et al*., 2017), with recommended k-mer size of 21 (jellyfish count -m 21 -o fastq.counts -C -s 500000000 - t 5 Genome_R1.fastq GenomeR2.fastq).

### Genotyping and linkage map construction

A cross between the sexual and asexual strains was set up to construct a linkage map for QTL mapping (Figure 1a). Virgin sexual AO females were individually crossed to asexual KG males (n=20) and provided with 100 hosts for oviposition (Figure 1a). After approximately two weeks, wasp pupae were isolated to collect virgin F1 females (n=23) that were again offered 100 hosts for oviposition. Recombinant F2 (haploid) males (n=188) were collected and stored at −80°C until DNA extraction and genotyping. DNA was extracted using a modified high salt protocol (adjusted from Aljanabi & Martinez, 1997). In short, each wasp was homogenized in a 1.5*ml* Eppendorf tube with homogenization buffer (0.4*M* NaCl, 10*mM* Tris:HCl pH8.0, 2*mM* EDTA) using a plastic pestle. After adding proteinase K, the tissue was incubated overnight at 55°C. After isopropanol precipitation, the DNA was dissolved in 20 *μl* MQ water.

The 188 F2 males were genotyped for the 113 SNPs using the high-throughput Kompetitive Allele-Specific PCR robot genotyping system (KASP, Semagn, Babu, Hearne, & Olsen, 2014) at the Institute of Biology Leiden (The Netherlands). Two allele-specific forward primers with a single difference at the SNP nucleotide position, and one common reverse primer (Table S2), were designed for each SNP fragment sequence. Each forward primer is attached to a unique unlabeled tail sequence at the 5’ end (15bp) which can be detected by fluorescence (Semagn *et al*., 2014).

The initial genotyping data set was first cleaned up by removing SNPs with >50% missing genotypes, which resulted in 96 suitable SNPs for linkage mapping. MapDisto software (version 1.7.7; Lorieux, 2012) was used to construct the genetic linkage map using the “BC” (backcross) population type setting to accommodate the haplodiploid sex determination system and cross design (Figure 1a). The “autoripple” function was used to order the loci per linkage group. Kosambi’s mapping function (Kosambi, 1944) was used to translate recombination fractions into map distances. The linkage map was visualized with MapChart (version 2.2, Voorrips, 2002).

### QTL mapping

In total 221 BC2 and 32 BC4 introgressed females were genotyped for the 96 SNP markers using the KASP platform. DNA was extracted from homogenized whole bodies with a Qiagen DNeasy kit after overnight treatment with 10% proteinase K (Qiagen) at 56 °C. The samples were processed on a BioSprint 96 workstation (Qiagen) with standard BS 96DNA program, resulting in 200 *ul* Buffer AE (Qiagen) DNA elution.

QTL analyses were conducted in the R package R/qtl v 1.46-2 (Broman & Sen, 2009; Broman, Wu, Sen, & Churchill, 2003). The genotype data were converted to the backcross format, with homozygotes encoded as ‘AA’ and heterozygotes as ‘AB’ (Table S3) (Broman & Sen 2009). Duplicated markers were identified and removed with the ‘findDupMarkers’ function. After removing markers with no genotype data, 92 markers remained. For each G2 or G4 recombinant female (n=253 in total), the probability of the allelic state at every map position, conditional to the observed genotype for the segregating SNP markers, was estimated with a hidden Markov model, allowing for 0.001 genotyping error rate and missing genotype data. The genetic map was also re-estimated from the backcross by the Lander-Green algorithm (hidden Markov model) for further QTL analysis using R/qtl (Broman & Sen 2009, Figure S1. QTLs for the female functional virginity were identified by Haley–Knott regression (HK) or the expectation maximization (EM) algorithm (Haley & Knott, 1992). The G2 and G4 generations were analyzed jointly. Function ‘scanone’ was run first for a binary model using 1,000 permutations, and generation as a co-variate. Both methods detected a single QTL on linkage group 12 at position 7 cM or 8 cM, respectively. The 5% threshold of logarithm of the odds (LOD) in both methods was 2.4. The ‘find.marker’ function was then used to identify the closest markers to the QTL and calculate the phenotypic effects of the markers on either side of the QTL using the function ‘effectplot’. The 95% confidence interval of the QTL region was calculated with function ‘bayesint’. EM and HK models with the ‘scantwo’ function were also run, but did not provide evidence for interactions between linkage groups between two independent QTLs. Finally, a linear regression was applied to investigate the effect of generation and an interaction between generation and the QTL (see details in Result section), and to estimate the percent of phenotypic variance explained by the QTL.

### Genome assembly and annotation

We assembled the genome of the asexual KG strain of *A. japonica* using PacBio long-read Single-Molecule-Real-Time (SMRT) sequencing (BioProject: PRJNA661661, accession number: JADHZF000000000). Samples of 50 milligrams of asexual KG females (approximately 120 individuals) were used for DNA extractions with Genomic-tip 100/G (Qiagen) according to the manufacturer’s protocol. A single library was constructed with SMRTbell, and BluePippin was used to select the >10kb long fragments. The library was run on 30 PacBio RSII cells. In total 2,189,795 (post-filter) reads were obtained, representing an approximate coverage of 94X. PacBio reads were then *de novo* assembled with FALCON (v0.2.2) with default parameters for insect datasets, but with modifications to fit our study species (Chin *et al*., 2016), i.e. length_cutoff_pr = 12000, pa_HPCdaligner_option = -v -dal128 -t16 -e0.75 -M24 - l4800 -k18 -h480 -w8 -s100, pa_Dbsplit_option = -x500 -s400, falcon_sense_option = --output_multi -min_idt 0.70 -min_cov 5 -max_n_read 200 -n_core 8. The draft genome was then polished with Quiver (2.2.1) (Chin *et al*., 2013).The quality and summary statistics of the assembled genome were analyzed with QUAST v4.6.3 (Gurevich, Saveliev, Vyahhi, & Tesler, 2013). BUSCO3 was used to estimate the completeness of the assembled PacBio KG genome (Simão, Waterhouse, Ioannidis, Kriventseva, & Zdobnov, 2015).

RNAseq datasets originally generated for a different study were used to annotate the KG genome. These datasets were obtained from ovary tissues of both sexual and asexual *A. japonica* strains. Long-read SMRT sequencing of RNA was conducted on 100 pooled ovaries of asexual KG females, and ovaries from sexual females of the population IR (Iriomote-jima island Japan, Murata *et al*., 2009). Dissected ovaries were disrupted in Trizol (Invitrogen, Carlsbad, CA, USA) and transferred to phase lock gel tubes. 200 μL chloroform was added to the 1 mL Trizol solution, shaken for 15 s and centrifuged for 10 min at 12.000 g at 4°C. The resulting upper layer was transferred to the gDNA Eliminator spin columns of the RNeasy Plus Micro Kit (Qiagen) and further RNA extraction proceedings followed the kit protocol. The Teloprime (Lexogen) method was used to synthesize and amplify full-length cDNA. The library was constructed with SMRTbell and sequenced on two cells of the PacBio Sequel. High-quality circular consensus sequences were used to identify full-length mRNA sequences, based on the presence of the primers and polyA sequences. IsoSeq pipeline (v3) was then used to perform the analysis and polish isoform sequences (*Gordon et al., 2015*). The high-quality isoform sequences were subsequently aligned with GMAP against the PacBio genome assembly of the KG strain (Wu & Watanabe, 2005). Finally, the generated transcriptome was functionally annotated with Blast2go v4.1 (Götz *et al*., 2008).

### Candidate gene analysis

Our QTL mapping approach uncovered a large QTL peak spanning over at least 4.23 Mb of the *A. japonica* genome assembly, in two non-overlapping contigs (see results). To obtain a list of candidate genes in this region, we used the original annotations from the PacBio genome (based on *A. japonica* transcriptomes from ovary tissue), and improved these using publicly available transcriptomes of three parasitoid wasp species (*Diachasma alloeum, Fopius arisanus*, and *Microplitis demolitor*; NCBI BioProject accession numbers: PRJNA284396, PRJNA259570, PRJNA214515 respectively) to ensure genes not expressed in ovaries were included. We were able to annotate 131 candidate genes in the QTL region (see results).

We hypothesized that female resistance to mating may be a female-specific trait and be regulated by genes featuring sex-biased expression. We screened the 131 candidate genes for sex-biased expression in a dataset of the sexual sister species *A. tabida*, which was originally generated for a different purpose. The *A. tabida* data set was obtained by extracting RNA from whole-bodies of male and female *A. tabida*, with TriZol (Invitrogen, Carlsbad, CA, USA) according to manufacturer’s protocol. Individual RNA extractions of 11 females were pooled for one female library, and of 25 males for one male library. RNA libraries were prepared and sequenced on the Illumina HiSeq2000 at the UMCG Sequencing Facility. In total, 20.2 million and 26.6 million reads were retained in the female and male libraries respectively (BioProject: PRJNA661661,SAMN16067724, SAMN16067725), after the trimming step to remove adaptors and low quality reads using Trimmomatic v0.33 with default parameters (Bolger, Lohse, & Usadel, 2014).

We performed *de novo* transcriptome assembly with Trinity v 2.4.0 using default parameters (Haas *et al*., 2013), and performed differential gene expression analysis between sexes with the R package edgeR v3.4 (Robinson *et al*. 2010; McCarthy *et al*. 2012; Chen *et al*. 2015). The details of differential expression analysis was described previously (Ma *et al*. 2018a, 2018b). Briefly, the trimmed reads of the female and male sample were mapped to the assembled transcriptome with Kallisto v.0.43.0 (Bray, Pimentel, Melsted, & Pachter, 2016). Read counts of the output from Kallisto mapping were imported for gene expression analysis in edgeR v3.4 (McCarthy *et al*., 2012; Robinson *et al*., 2010) and filtered with average Log_2_(CPM) > 0 per sample, followed by normalizing the expression by trimmed mean M values (TMM). Normalized expression counts for each sample were used to calculate sex bias using standard measures and Benjamini-Hochberg correction for multiple-testing with false discovery rate (FDR) of 5%. As we did not have biological replicates, the recommended dispersion value 0.4 was used in edgeR (Chen *et al*. 2016), and stringent criteria were applied when calling sex bias (|log_2_FC| ≥2, or fold change ≥4).

One candidate gene out of 15 with different expression between the sexes in *A. tabida*, *hormone receptor 4*, was particularly interesting due to this gene function association with mating propensity. We therefore examined this gene for putative loss-of-function mutations in the asexual *A. japonica* strain. To compare the gene sequences between sexual and asexual strains of *A. japonica*, we mapped the Illumina short reads of the sexual AO strain against the PacBio genome using bwa mem with default settings (Li & Durbin, 2009). Before and after read filtering, read mapping quality was assessed with Samtools (v1.3) flagstat function (Li et al., 2009), and filtered by quality threshold >20. SNP calling was conducted on the mapped SAM files with BCFtools (v1.7, Narasimhan et al. 2016), and mapped reads with mapping depth coverage below 3 and above 80 were filtered out. Final SNPs were then called with vcfutils.pl scripts from BCFtools (v1.7) and plotted with a 500bp sliding window for the two QTL contigs using ggplot2 in R (v3.6.3, R Core Team 2017). We also compared *hormone receptor 4* sequences of 34 additional hymenopteran species, by identifying *hormone receptor 4* orthologs through BLAST against the NCBI database, and generating multiple alignments of the detected orthologs using Geneious v8.1.7 (https://www.geneious.com).

## Results

### Female functional virginity

We previously documented that asexual females of *A. japonica* rejected all mating attempts when confronted with males (Ma *et al*. 2014a). We classified this female resistance to mating as representing female functional virginity (sensu (Huigens & Stouthamer, 2003)). To determine the genetic basis of female functional virginity, we previously introgressed alleles from the asexual KG strain into the sexual AO strain for four consecutive generations (summarized in Table 1, see also Ma *et al*. 2014a). In the present study, females from these same introgression generations were used for QTL analysis.

**TABLE 1.** Females rejecting matings over successive generations of introgression of the asexual genome into the sexual genome in *Asobara japonica*. Sample sizes, number and proportion of successful and failed matings for four consecutive generations (*Ma et al*. 2014a, b).

| Generation | Percentage of asexual genome | Sample size | No. of successful matings | No. of failed matings | Percentage of successful matings |
| --- | --- | --- | --- | --- | --- |
| G0<br>(sexual strain) | 0% | 56 | 50 | 6 | 89.29% |
| F1 | 50% | 49 | 46 | 3 | 93.88% |
| G2 | 75% | 225 | 121 | 104 | 53.78% |
| G3 | 87.50% | NA | NA | NA | NA |
| G4 | 93.75% | 51 | 21 | 30 | 41.18% |
| G0<br>(asexual strain) | 100% | 20 | 0 | 20 | 0.00% |

### Linkage map construction

The DIAL pipeline initially identified 21,976 SNPs, and a series of filtering steps led to 420 SNPs for further testing and selection (see Methods for details). Eventually, a set of 96 SNPs was used to construct a linkage map from the genotypes of 188 F2 haploid males (Figure 2). Linkage groups were determined with a minimum LOD score of 3 and a maximum recombination frequency of 0.35 between the linked pairs of SNP markers. After this first mapping, the double recombinants were used to identify potential genotyping errors with an error detection *P*<0.05. Using this threshold, 25 erroneous candidates were removed which were coded as missing data. Next, Chisquare tests were used to check for segregation distortion from Mendelian expectations (1:1 for haploid F2 males). Segregation distortion was observed for 13.5% of SNP markers (13 out of 96, Chi-square test, *P*<0.05), nine SNPs were skewed to the asexual genome and four to the sexual genome. Interestingly, one linkage group (LG7) consisted of seven markers that all significantly deviated from Mendelian expectations, with the asexual genome variant being more common than expected. Four of the seven markers with the most extreme segregation distortion (*P*<0.000001) were removed from LG7 to enable proper QTL analysis (Table S4). Segregation distortion could be due to genetic incompatibility between strains or some other factors that remain to be investigated. The final map consisted of 16 linkage groups, and the total length was 667.32 cM with an average distance between markers of 7.3 cM (Figure 2).

**FIG. 2.**
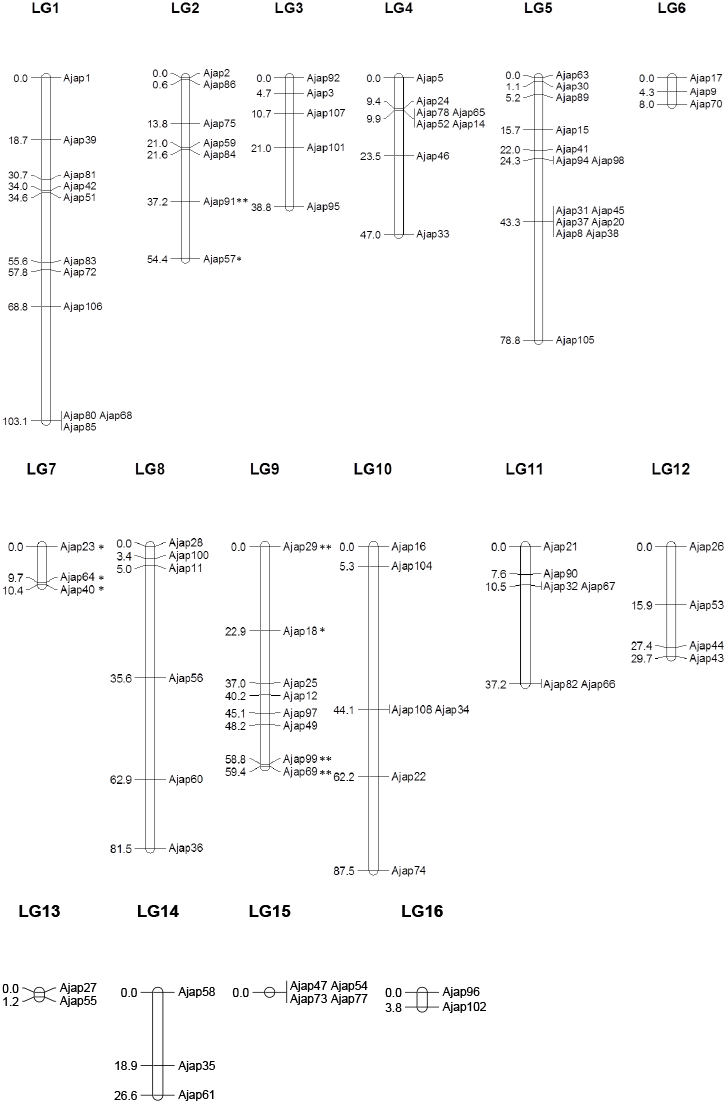
Genetic linkage map of *Asobara japonica*. For each linkage group, the relative position (centimorgans) is indicated on the left side and the SNP markers are shown on the right side. ** denotes *P* <0.01, * denotes *P* <0.05 in chi-square tests for segregation distortion. LG=linkage group.

### QTL of asexual female functional virginity

In total, 253 sex-asex introgressed females (221 G2 and 32 G4) were genotyped (Figure 1, Table 1). QTL analysis identified a single QTL explaining 48.8% of the variation among females for the presence or absence of functional virginity (Table 2, Figure 3). It was located on LG12 at 7 cM (based on the EM algorithm) or 8 cM (on the HK regression), with very high LOD values (41.3 and 40.8 respectively). No other significant QTL was detected across the genome above the 5% LOD threshold of >2.51, estimated from 1,000 permutations. Furthermore, no significant interactions of paired markers between linkage groups were detected, neither by the scantwo function nor by the multiple QTL function in R/QTL. The 95% confidence interval of the QTL location ranged between 4-10 cM with the EM method and 6-11 cM with the HK method (Figure 3). The generation (BC2 or BC4) was not a significant variable in the QTL model.

**FIG. 3.**
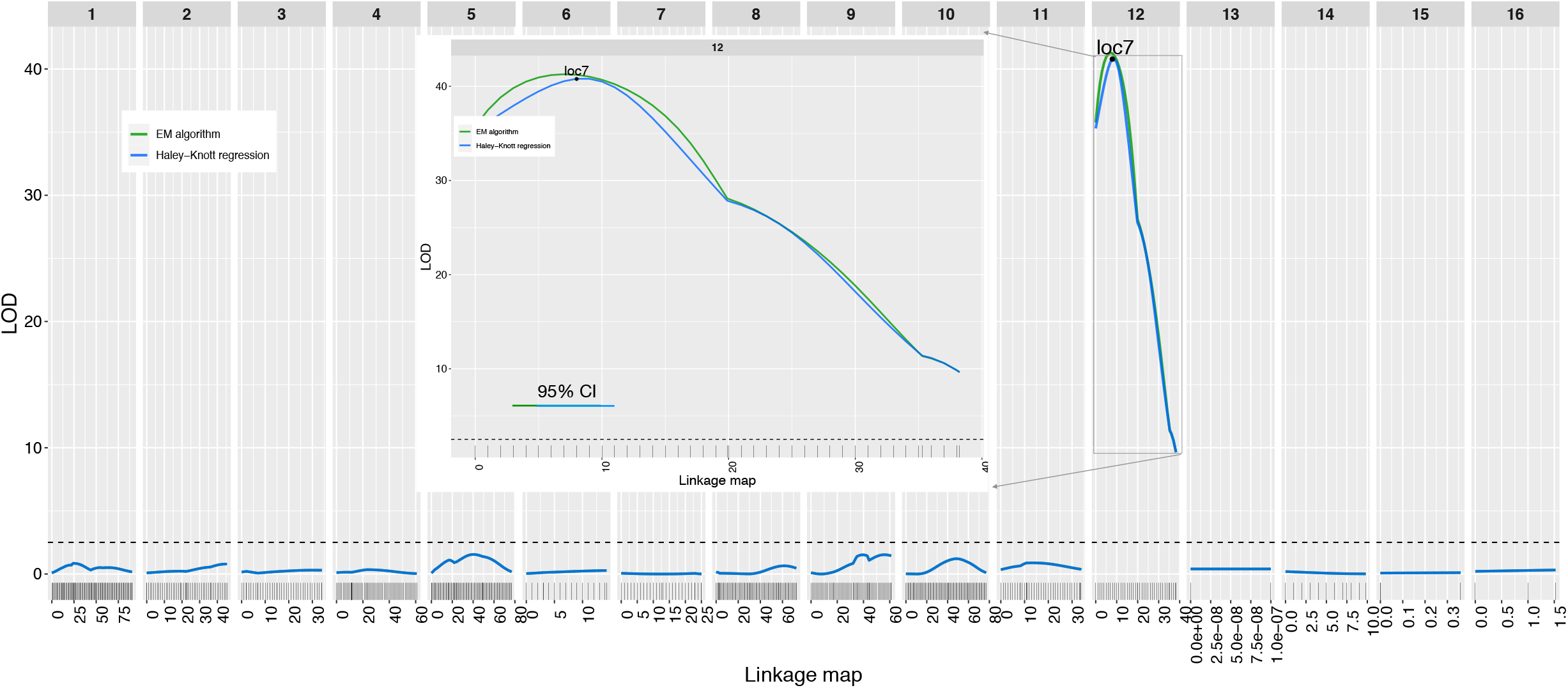
A single QTL for female functional virginity in *A. japonica*. The horizontal line corresponds to the 5% genome-wide significant threshold from permutation tests. The inserted figure shows 95% of confidence interval for the detected QTL at LG12, with range of 4-10 cM for expectation maximization (EM) and 6-11 cM for Haley-Knott regression (HK) method. LG=linkage group.

**TABLE 2.** Genome-wide significant QTLs for female mating success (sex-asex introgressed females) in *A. japonica*. Binary model formula: y ~ Generation + Q1 + Generation:Q1, generation and the interaction with Q1 were not significant.

| Variable | Linkage group | Location (cM) | Variation explained (%) | LOD score | P value |
| --- | --- | --- | --- | --- | --- |
| Model | NA | NA | 48.75 | 36.73 | 0 |
| QTL | 12 | 6 | 48.75 | 36.73 | <2e-16 |
| Generation | NA | NA | 0 | 0 | 1 |
| Generation:<br>QTL | NA | NA | 0 | 0 | 1 |

Two markers, Ajap26 (7 – 8 cM away from the QTL peak) and Ajap53 (7.9 – 8.9 cM away, Figure 3), delineate the QTL 95% confidence intervals (4-11 cM from EM and HK model estimate). Females produced daughters (>80%) when the markers were heterozygous (C:G for Ajap26 and A:T for Ajap53), and only sons when they were homozygous (G:G and A:A respectively, which are the genotypes in the asexual strain; Figure 4). Note that, given the cross scheme (sex-asex introgressed females always mated to asexual males), the alternative homozygote C:C or T:T (fixed for the sexual strain) did not occur in the sex-asex introgressed females. These results corroborate the prediction of a single recessive locus to be the determinant of female functional virginity (Ma *et al*. 2014a).

**FIG. 4.**
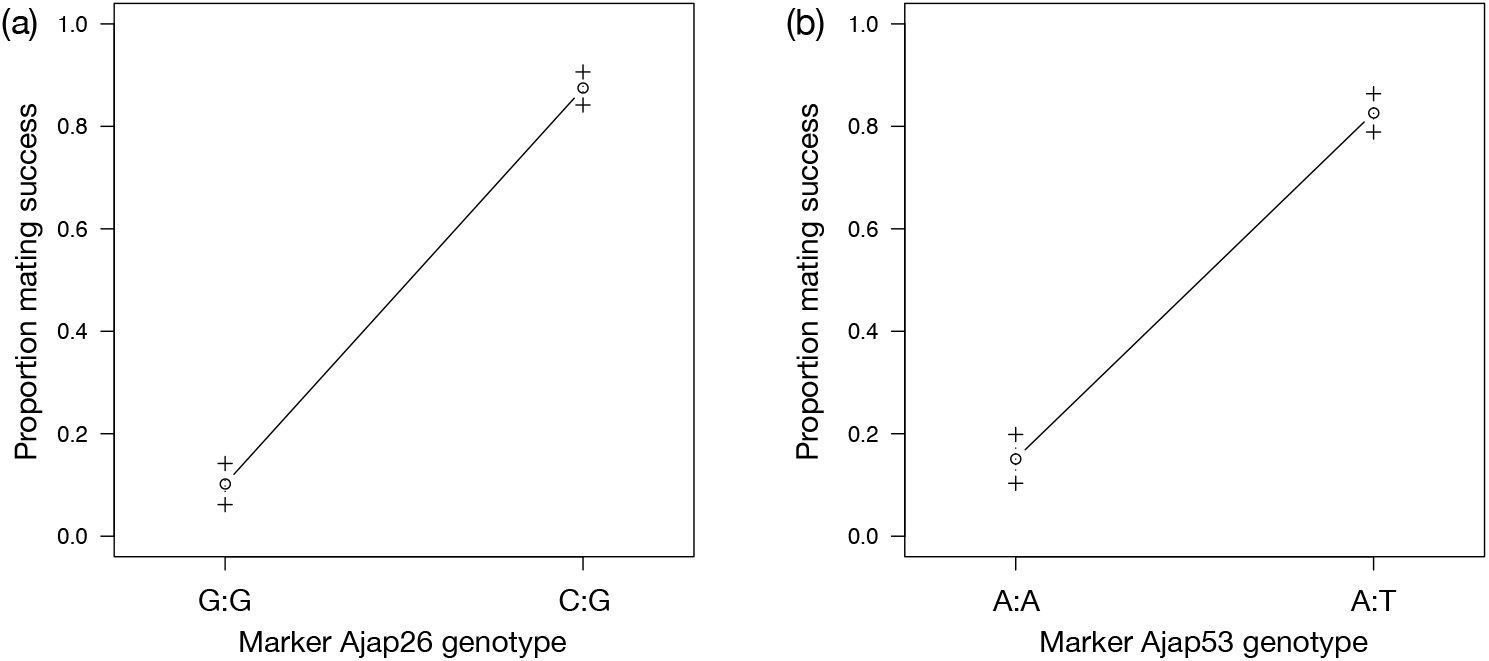
Genotype and associated proportion of successful mating for the introgressed sex-asex females at each of the two most closely linked markers flanking the detected QTL. Marker Ajap26 (a) and marker Ajap53 (b).

### Candidate gene analysis

The assembled genome length is 271.51Mb, suggesting that our PacBio genome assembly is 89.3-96.4% complete (see Table 3 for additional statistics). Furthermore, BUSCO v3.0.1 identified 97% complete (C), 0.5% fragmented (F), and 96.4% singlecopy (S) genes. BUSCOs using the Insecta dataset (n=1658, C: 97.0% [S:96.4%, D (duplicated):0.6%], F:0.5%, M (missing):2.5%, n:1658). The assembled transcriptome of *A. tabida* was composed of 34,140 transcripts longer than 300bp, and 2,810 (8.2%) transcripts were differentially expressed between the sexes with stringent criteria of |log_2_FC| ≥2 (Table S5).

**TABLE 3.**
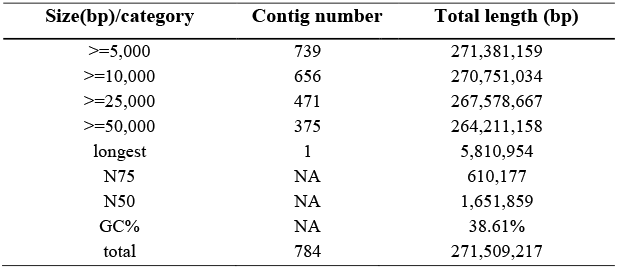
Summary of the assembled PacBio genome of the asexual KG female of *A. japonica*.

The two SNP markers that identified the QTL mapped to two non-overlapping contigs of the KG PacBio genome, contig0058 (1.78Mb) and contig0035 (2.45Mb). Based on the total genetic map length of 667.43cM, assuming even marker density and using an estimated genome size 281.6 - 304.1 Mb, the two contigs comprise 58-63% of the genomic region under the QTL peak. As female functional virginity trait (i.e. female resistance to mating) is likely a female-specific trait, we speculated that genes underlying its regulation may show sex-biased expression in sexual species.

The two contigs associated with the QTL comprised 131 genes annotated with transcriptomes from different wasp species (Table S6). Among the 131 genes, 15 genes showed sex-biased expression in whole-body RNA libraries of the sexual species *A. tabida* (Table 4). We studied one interesting gene in more detail, *hormonal receptor 4*, with significant female bias (Log_2_(male/female) = −3.56). The coding sequence of *hormone receptor 4* from *Diachasma alloeum* was used to detect the exon and intron boundaries in the (asexual) *A. japonica* genome. A premature stop codon was detected in the asexual KG strain, however, a nonsense mutation was also present in the sexual AM strain. This nonsense mutation was also detected in several other sexual hymenopteran species (Figure S2, File S1). A relatively higher density of SNPs between sexual and asexual *A. japonica* strains was found in the promoter than intron regions (upstream up to 5kb of the start codon), and a lower density of SNPs was detected in the exons compared to promoter and intron regions (Figure S3). These results are consistent with the idea that mutations in regulatory regions, and/or SNPs could potentially lead to differences in expression or function of this gene in *A. japonica*.

**TABLE 4.** The 15 candidate genes with sex-biased expression in the sexual species *Asobara tabida*. All but one show female-biased expression. in bold indicates shared putative candidate gene with mutations in asexual genome of *Leptopilina clavipes* (Kraaijeveld *et al*., 2016), # in bold indicates shared putative candidate gene with differential expression between sexual and asexual stick insects in *Timema* (Parker *et al*., 2019).

| Species | Gene /Transcript | Gene function | Sex bias | Log2(male/female) |
| --- | --- | --- | --- | --- |
| *A. japonica* | transcript/15564 | **CD151 antigen**# | male | 3.98 |
| *A. japonica* | transcript/543 | DNA replication ATP-dependent helicase/nuclease DNA2 | female | -3.62 |
| *A. japonica* | transcript/11757 | PREDICTED: uncharacterized protein LOC105264533 | female | -4.13 |
| *A. japonica* | transcript/349 | cordon-bleu protein-like 1 | female | -4.2 |
| *A. japonica* | transcript/4407 | **nuclear pore complex protein Nup50** * | female | -11.04 |
| *A. japonica* | transcript/7189 | protein lin-9 homolog | female | -3.02 |
| *A. japonica* | transcript/1434 | putative ankyrin repeat domain-containing protein 31 | female | -3.88 |
| *A. japonica* | transcript/15419 | ubiquitin-conjugating enzyme E2 L3 | female | -3.62 |
| *A. japonica* | transcript/16014 | vesicle-trafficking protein SEC22b-B | female | -7.23 |
| *D. alloeum* | XM_015253230.1 | homeobox protein Nkx-6.2 | female | -3.6 |
| *D. alloeum* | XM_015260178.2 | unconventional myosin-XIX | female | -4.07 |
| *D. alloeum* | XM_015262299.1 | phospholipase D1 | female | -10.43 |
| *D. alloeum* | XM_029125918.1 | uncharacterized LOC107035734 | female | -4.2 |
| *F. arisanus* | XM_011307786.1 | hormone receptor 4 | female | -3.56 |
| *F. arisanus* | XM_011317030.1 | open rectifier potassium channel protein 1 | female | -4.85 |

## Discussion

Our aim was to investigate the genetic basis of female functional virginity in *A. japonica*. This required, as a first step, to generate a linkage map and genomic datasets that will be useful for comparative genomic studies across hymenopteran species. With an average inter-marker distance of 7.3 cM, the *A. japonica* map has a comparable resolution to maps for other hymenopterans, including 4.0 cM for the bumblebee *Bombus terrestris* (Stolle *et al*., 2011), 7.5 cM for the honeybee *Apis mellifera* (Solignac *et al*., 2004), 11.1 cM for *Nasonia vitripennis* (Pannebakker, Watt, Knott, West, & Shuker, 2011), 17.0 cM for *Bracon hebetor* (Antolin *et al*., 1996) and 17.7 cM for *Trichogramma brassicae* (Laurent *et al*., 1998). Moreover, based on the estimated genome size of 281-304 Mb, the recombination rate of *A. japonica* is 2.20~2.37 cM/Mb, compared to ~1.5 cM/Mb for *Nasonia* (Pannebakker, Watt, Knott, West, & Shuker, 2011), ~4.76 cM/Mb for *Bombus terrestris* (Stolle *et al*., 2011) and ~22.8 cM/Mb for *Apis mellifera* (Solignac *et al*., 2004). This, and the fact that the 16 linkage groups defined by the SNP map agree with earlier estimates of the number of chromosomes (12-17 chromosomes based on cytological data; Gokhman 2009; personal communication G. Massimo), indicate the accuracy of the constructed linkage map.

Our QTL analysis revealed that female functional virginity is associated with a single QTL with major effect. This is consistent with the previous prediction of a single recessive locus based on the segregation of phenotypes in our introgression lines (Ma *et al*. 2014a). In this previous study, *Wolbachia*-cured asexual females of *A. japonica* were found to have reduced attractiveness to males (only 55% asexual females attracted sexual males vs. 100% sexual females), and a complete loss of mating acceptance resulting in a total absence of matings. As a result, these females produced only haploid sons (Ma *et al*. 2014a). A simple genetic architecture for female functional virginity was also found in two other studies of wasps with endosymbiont-induced asexuality. Inability to fertilize eggs segregated as expected for one or few loci with recessive effects in *Telenomus nawai*, and as expected for few loci with dominant effects in *Trichogramma pretiosum* with additional minor-effect modifiers (Jeong and Stouthamer, 2005; Russell and Stouthamer, 2011).

Our QTL analysis identified a single large-effect QTL on LG12 at 7 or 8 cM (95% CI: 4-11 cM), strongly associated with female functional virginity. Annotation of the QTL region identified 131 physically-linked candidate genes. To our knowledge, this is the first study to identify candidate genes for female functional virginity, a trait that is predicted to evolve in the transition from sexuality to asexuality (Huigens & Stouthamer 2003). Because female functional virginity is female-specific, we hypothesized that the underlying gene(s) would show differential expression between sexes in sexual species. In total, 15 of the 131 genes were sex-biased in *A. tabida*, the sexual sister of *A. japonica*, and all but one had female-biased expression. The molecular changes in genes associated with the decay of sexual traits are still poorly known. A review (Arbuthnott, 2009) on the genetic architecture of sexual traits influencing premating isolation, such as behavioral signaling and courtship traits, found 69% (25 of 36) to be encoded by few loci with relatively large effects, and suggested changes in courtship behavior may often evolve or decay quickly. Little is still known about the genetic basis of female receptivity, but auditory and/or olfactory receptors are likely candidates (Laturney & Moehring, 2012). One gene, *hormone receptor 4*, is particularly interesting in this respect, because this gene functions as a steroid hormone receptor, and may therefore be relevant to mating resistance. Furthermore, a recent study also showed that this gene regulates the mating propensity in the malaria mosquito *Anopheles gambiae* (Gabrieli *et al*. 2014). We detected a premature stop codon in this gene in both the sexual and asexual strains, as well as in several other hymenopteran species. Moreover, we found multiple SNPs in the upstream (putative promoter) region, and in the gene itself, differentiating the sexual and asexual *A. japonica* strains. Further functional analyses would be required to confirm the role of this gene in female functional virginity in *A. japonica*.

Few studies have identified genes that are subject to change following transitions to asexuality. Kraaijeveld et al. (2016) compared genomes of sexual and asexual linages of the parasitoid wasp *Leptopilina clavipes*, and identified 16 genes with deleterious mutations (including frame shifts and early stop codons) associated with asexuality. One of our female virginity candidate genes, *nuclear pore complex protein Nup50*, was also among these 16 genes of *L. clavipes. Nup50* is involved in mRNA transport, or positive regulation of transcription by RNA polymerase II and predominantly interacts with transcriptionally active genes inside the nucleoplasm, in particular those involved in developmental regulation and the cell cycle (Buchwalter, Liang, & Hetzer, 2014; Kalverda, Pickersgill, Shloma, & Fornerod, 2010). In future studies, a near chromosomal-level genome assembly to completely cover the QTL region would be helpful to evaluate whether there are more candidate genes for female functional virginity in *A. japonica*. Meanwhile, functional analyses of the identified 131 candidate genes, especially the 15 ones with sex-biased expression, can be undertaken to further elucidate the molecular basis of female functional virginity in this species. Whether certain genes are particularly affected by a shift to asexuality requires more comparative studies on a broader array of species.

Sexual traits, including female mating behaviours, are costly, which may lead to such traits being selected against under asexuality (King & Hurst, 2010; Russell & Stouthamer, 2011; Stouthamer *et al*., 2010). Theoretical modeling of the spread of asexuality-inducing endosymbionts in mixed populations with uninfected and infected females, has confirmed that female functional virginity is favored under sex-ratio selection (Stouthamer *et al*., 2010). Such selection might be stronger in haplodiploid species because unmated females will produce only haploid male progeny that balances the sex ratio. Once a mutation for rejection to mate has swept through an asexual population, the region comprising the genes associated with mating resistance does not get smaller due to lack of recombination in asexuals. This may explain the large size of the single QTL detected in our study. Overall, our results are consistent with mutation(s) in the genomic region consisting of a single gene or a cluster of linked genes underlying rapid evolution of female functional virginity in the transition to asexuality.

## Acknowledgements

We thank Rogier Houwerzijl and Peter Hes for assistance with wasp culturing, Ken Kraaijeveld, Barbara Reumer and Fabrice Favre for supplying *A. japonica* strains. We thank Marloes van Leussen for the PacBio DNA/RNA extractions, Fangying Chen for the wasp dissections and Klaas Vrieling for KASP analyses support. We thank Mathijs Nas for performing the verification test of the *in-silico* SNP markers, Bob Dröge and Aakrosh Ratan for assistance with the DIAL pipeline, and Bregje Wertheim for valuable discussions. We thank the DroParCon consortium (https://wiki.gcc.rug.nl/wiki/DroparconStart) for providing Illumina short reads of both strains of *A. japonica*, and this consortium was supported by Van Gogh grant to B. Pannebakker and Fabrice Vavre. W.-J. Ma was supported by Ubbo Emmius Fellowship from the University of Groningen. X. Li is supported by China Scholarship Council (CSC) Scholarship no. 201606330077. This work was supported by grants no. 854.10.001 and no. 824.15.015 from the Netherlands Organization for Scientific Research (NWO).

## Author Contributions

**Conceptualization**: Wen-Juan Ma, Bart Pannebakker, Tanja Schwander, Leo W.

Beukeboom, Louis van de Zande

**Data curation**: Wen-Juan Ma, Xuan Li, Elzemiek Geuverink, Seyed Yahya Anvar

**Genome, transcriptome assembly and annotation**: Xuan Li, Elzemiek Geuverink, Seyed Yahya Anvar

**Data analyses**: Wen-Juan Ma, Paris Veltsos

**Funding acquisition**: Leo W. Beukeboom, Louis van de Zande, Tanja Schwander

**Supervision**: Bart Pannebakker, Tanja Schwander, Louis van de Zande, Leo W. Beukeboom

**Visualization**: Wen-Juan Ma, Paris Veltsos

**Writing – original draft**: Wen-Juan Ma

**Writing – review & editing**: Wen-Juan Ma, Paris Veltsos, Leo W. Beukeboom, Tanja Schwander, Louis van de Zande, Bart Pannebakker, Elzemiek Geuverink, Xuan Li, Seyed Yahya Anvar

## Data accessibility

The data presented in this study can be accessed on NCBI Bioproject (PRJNA661661) with Accession No. (SRR12618696-SRR12618699, JADHZF000000000, SAMN16067724, SAMN16067725). All scripts for the analyses are deposited on GitHub: https://github.com/Wen-Juan/Decaytrait_qtl. All datasets and scripts will be released upon acceptance of this manuscript.

## Supporting information

**FIG. S1** Comparison of original linkage map (left) and adjusted map (right) following the hidden Markov model.

**FIG. S2** Multiple alignment of coding sequence of the candidate gene *hormonal receptor 4* of *A. japonica* and orthologs from 34 species (96 isoform sequences) in ants, bees and wasps. Black arrow on position 2120 indicates the SNP (G) causing early stop codon in some of these species.

**FIG. S3** SNP density plot per 100bp window of the candidate gene *hormonal receptor 4*, reads of sexual strain AO were mapped against the asexual KG female genomic regions of this gene. The upstream 5 kb starting from first nucleotide of start code is indicated in orange, and the green bars highlight exons.

**Table S1** List of primer pairs for verifying the in-silico generated SNP markers.

**Table S2** List of primer pairs for the SNP cluster sequences. For each primer pair, there are two forward primers named alleleX_forward and alleleY_forward, which is designed to distinguish the SNP position for each allele. There is a common reverse primer for each allele.

**Table S3** The converted genotypes of all 253 sex-asex admixed females for each of the 96 SNP markers used for QTL analysis, which were positioned on 16 constructed linkage groups.

**Table S4** The segregation distorted SNP markers and the Chi-square tests *P* values and significance (*P*<0.05).

**Table S5** Sex-biased gene expression numbers detected based on various Log_2_ fold change values.

**Table S6** All 131 annotated genes from four transcriptomic datasets, 37 from *Diachasma alloeum*, 46 from *Fopius arisanus*, 3 from *Microplitis demolitor* and 78 from *Asobara japonica*. Note that 33 annotations are for orthologous gene sequences in two or more species, these are merged in the list of 131 genes.

**File S1** The alignment of *hormonal receptor 4* coding sequence of *Asobara japonica* and additional 34 species from ants, bees and wasps.

