## Supplementary figures and images for "A single QTL with large effect is associated with female functional virginity in an asexual parasitoid wasp"

### Supplemental Figure 1

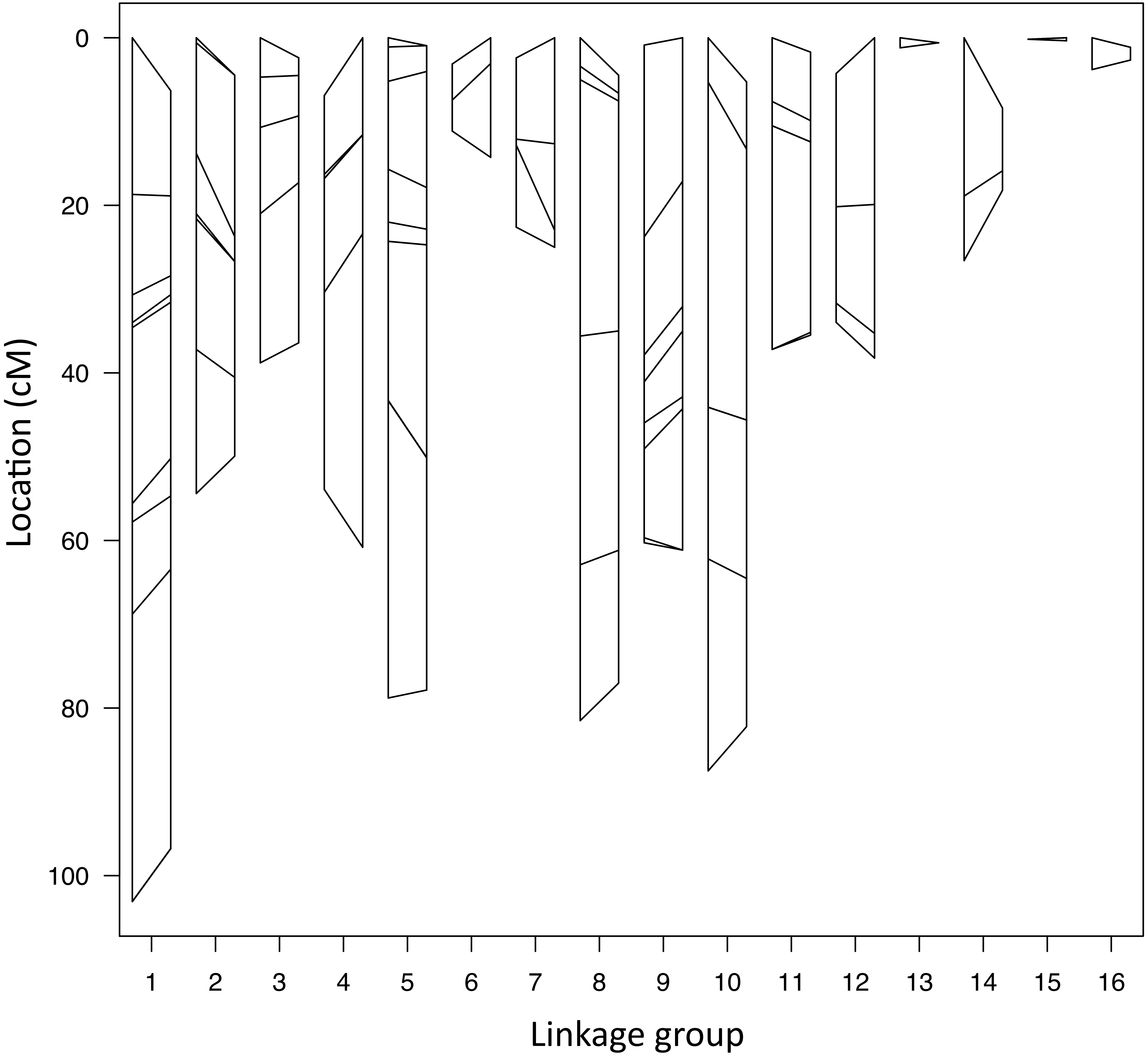

### Supplemental Figure 2

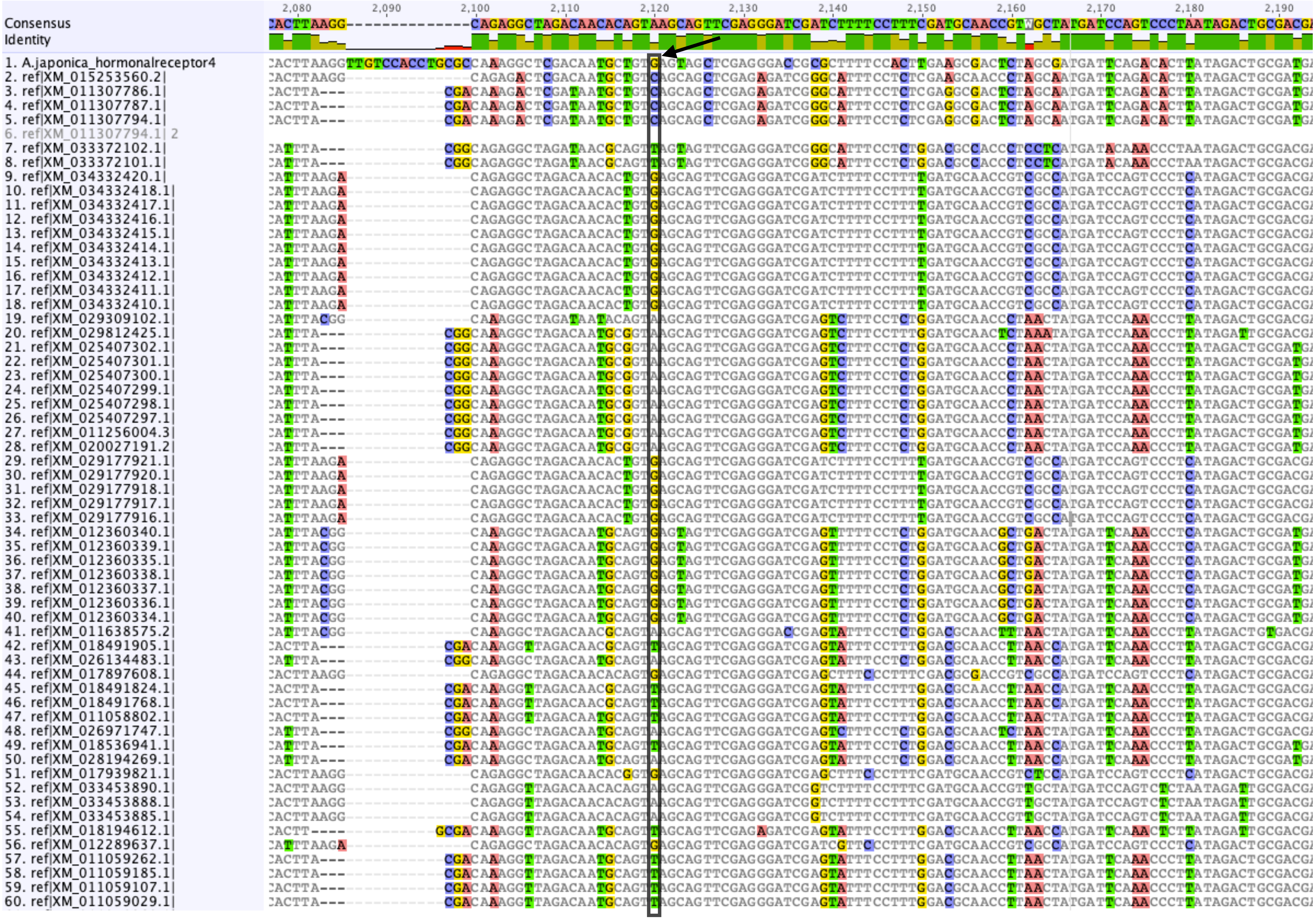


to be continued

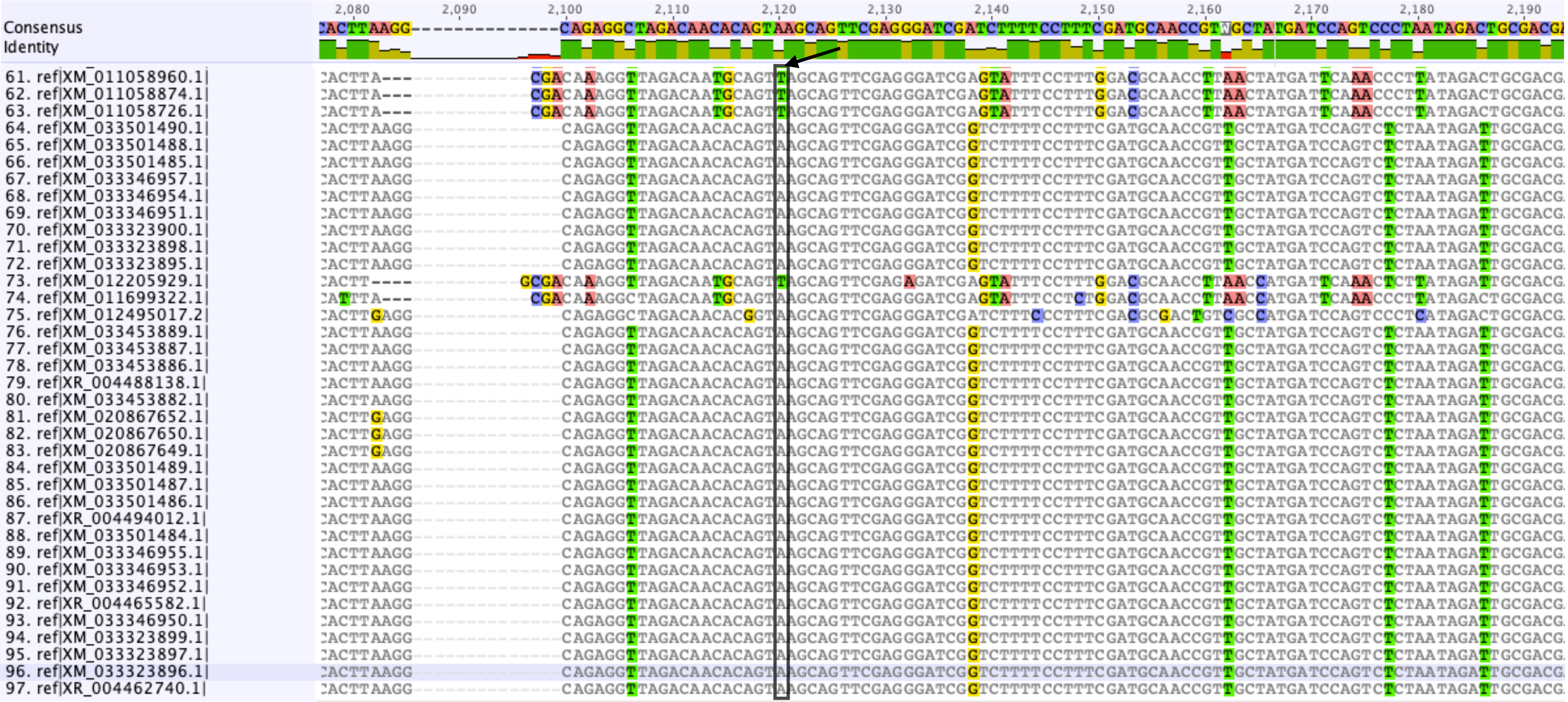

### Supplemental Figure 3

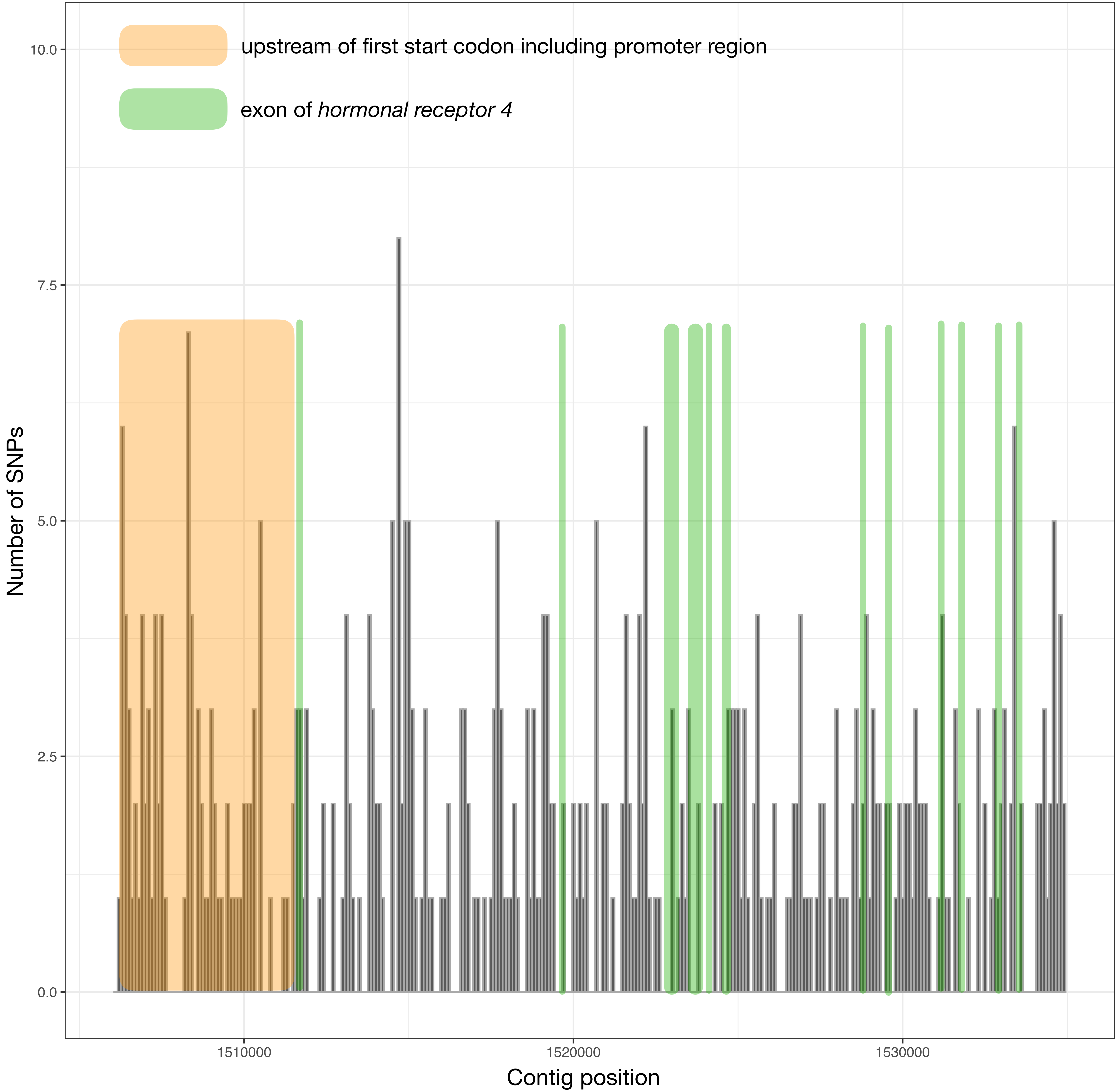
